# High-severity fires undermine resilience of black spruce-dominated boreal forests in eastern North America

**DOI:** 10.1101/2025.05.17.654490

**Authors:** Stelsa Fortin, Yan Boucher, Yves Bergeron, Martin Simard, Dominique Arseneault, Hugo Asselin, Martin Barrette, Victor Danneyrolles, Sylvie Gauthier, Francois Girard, Martin Girardin, Marc-André Parisien, Nelson Thiffault, Osvaldo Valeria

**Author notes:** Correspondence: Stelsa Fortin; Yan Boucher.

## Abstract

Climate-induced fire regime shifts may reduce post-fire resilience of black spruce-dominated (BS; *Picea mariana*) North American boreal forests. While post-fire vulnerability of immature BS stands has been extensively studied, no study has evaluated simultaneous effects of fire severity and seasonality on the post-fire regeneration of mature (> 60-year-old) BS stands. This study aims to quantify post-fire regeneration levels of BS and co-occurring tree species to assess ecosystem recovery and possible loss of resilience due to regeneration failure. We analyzed effects of seed bank conditions, fire regime characteristics (fire severity and seasonality), and seedbed conditions on BS post-fire regeneration in mature forests in Quebec, Canada. Post-fire regeneration density was extensively surveyed across ∼50 400 km^2^ through a network of 536 plots that were distributed in 21 fires, which burned between 1995 and 2016. One-third of plots failed to regenerate (< 1750 conifer seedlings/ha) at levels adequate to produce closed-crown forest, whereas one-fifth experienced compositional changes, mainly towards jack pine (JP; *Pinus banksiana*) dominance. Pre-fire basal area of BS and living *Sphagnum* ground cover increased BS post-fire regeneration, whereas high-severity crown fires and spring fires reduced it. These findings suggest that mature BS-dominated forests may lose resilience in response to high-severity and spring fires. Given the projected increase in fire severity, and the extension towards an early-fire season in response to climate change, our study suggests that post-fire regeneration failure may become more frequent over the coming decades, with potential negative consequences on ecosystem services that are provided by BS-dominated boreal forests.

## 1. Introduction

The North American boreal forest is characterized by a stand-replacing wildfire regime (Frelich et al. 2024), which influences vegetation dynamics by favouring species with regeneration traits that are adapted to fire (Keeley and Pausas 2022; Ruggirello et al. 2023). Although vegetation has responded to climate change throughout the Holocene (the past ca. 11,700 years) (Girardin et al. 2013, 2024; Aakala et al. 2023), disturbance-induced ecosystem changes that have been observed over the last few decades are occurring at an increasing rate (Remy et al. 2017; Baltzer et al. 2021). Ongoing climate change is altering fire regimes across the circumpolar boreal forest (Lehtonen et al. 2016; Coogan et al. 2019; Feurdean et al. 2020; Wang 2024), where temperatures are rising at one of the fastest rates on Earth (IPCC 2023). An increase in the number of days with warm temperatures and vapour-pressure deficits that are conducive to fire spread is expected (Augustin et al. 2022; Foster et al. 2022; Whitman et al. 2022), especially during the spring. Fires that occur in the spring, just after snowmelt but prior to the growing season, can quickly spread due to high wind velocities, which are associated with the retreat of the Polar Front and the low moisture content of plant tissues (Tymstra et al. 2021; Parisien et al. 2023). In Canada, the length of the fire season and the number of spring fires have increased since the early 1950s (Albert-Green et al. 2013; Flannigan et al. 2013; Hanes et al. 2019; Ahmed and Hassan 2023). Moreover, the increased frequency of droughts and extreme fire weather that is associated with climate warming are thought to cause more extreme fire behaviour, including an increase in fire severity (Parks and Abatzoglou 2020; Gaboriau et al. 2022; Jain et al. 2022).

Fire regime characteristics interact with pre-fire forest attributes to control post-fire forest recovery (Remy et al. 2017; Davis et al. 2023; Gaboriau et al. 2023; Boulanger et al. 2025). These characteristics can be grouped into three broad categories: temporal (e.g., fire frequency, return interval, and seasonality — the period when fires occur), spatial (e.g., fire size and shape), and magnitude (e.g., fire intensity, which refers to energy released, and fire severity, which describes the effects of fire on vegetation and soils). Fire severity in the canopy versus on (and in) the ground must be distinguished in the boreal forest, given that the two are not necessarily correlated. While the canopy burns within minutes to hours, thick organic soils can smoulder for weeks to months, leading to prolonged combustion.

In eastern North America, most of the boreal forest is dominated by black spruce (*Picea mariana* [Mill.] B.S.P.), hereafter abbreviated as BS, a conifer species that is well adapted to fire, and which can establish across a broad range of climatic and biophysical conditions (Viereck and Johnston 1990; Johnson 1992). To ensure self-replacement after fire, BS depends upon a large aerial seed bank in semi-serotinous cones, the post-fire opening of which can be delayed for several years (Greene et al. 2013; Splawinski et al. 2022). In addition, BS can multiply vegetatively by layering during long fire return intervals (Keeley et al. 2011). BS reaches reproductive maturity after about 50 years, but seed production peaks between 100 and 200 years, depending upon site fertility and growing degree-days (Viglas et al. 2013; Van Bogaert et al. 2015; Splawinski et al. 2022). Most seeds are released and germinate during the first 2-3 years following fire (St-Pierre et al. 1992; Gutsell and Johnson 2002; Greene et al. 2013), and seed availability and viability during this short time window are a sine qua non condition to ensure BS regeneration (Viglas et al. 2013; Perrault-Hébert et al. 2017). For optimal establishment and growth, BS seeds should land on moist mineral soil (Thomas and Wien 1985; Charron and Greene 2002; Wang and Kemball 2005). Unburned patches of *Sphagnum* moss may also serve as good germination beds, despite being suboptimal substrates for tree growth. Yet, their high-water retention capacity ensures seedling establishment and survival (Lavoie et al. 2007; Lloyd et al. 2007; Boiffin and Munson 2013; Mallik and Kayes 2018). In organic soils, high-severity fires tend to consume a thicker portion of the organic layer and are thus more likely to expose the mineral soil (Johnstone et al. 2009; Keeley 2008), but low-severity fires consume less of that layer, resulting in contrasting seedbed conditions (Lecomte et al. 2006; Veilleux-Nolin and Payette 2012). Although BS is adapted to high-severity crown fires, extreme severity may also be detrimental to species recruitment, which could lead to regeneration failure (Perrault et al. 2017; Reid et al. 2023). The species’ small seeds are enclosed in thin cone scales that are vulnerable to high-severity fires (Arseneault 2001; Splawinski et al. 2019b). BS regeneration failure can result in open canopy BS woodlands, in deciduous stands, or in stands that are dominated by jack pine (JP; *Pinus banksiana* Lamb.) (Hart et al. 2019; Baltzer et al. 2021; Burrell et al. 2021; Stevens-Rumann et al. 2022).

It is widely accepted that immature BS stands are at high risk of post-fire regeneration failure due to undeveloped seed banks (Girard et al. 2009; Viglas et al. 2013; Perrault-Hébert et al. 2017; Hart et al. 2019), but very little information is available regarding the vulnerability of mature BS stands to fire regime characteristics and the drivers that are involved in species post-fire recovery. Therefore, this study aimed to quantify the relative importance of various environmental variables affecting the post-fire regeneration of mature BS stands across a 50 400 km^2^ boreal landscape in eastern North America. We hypothesized that high-severity fires and unfavorable seedbed conditions would hamper post-fire regeneration and possibly result in regeneration failure. To verify this hypothesis, we established a large sampling network of 536 plots between the summers of 2017 and 2020. The plots were located within sites where 21 fires had burned between 1995 and 2016. The sampled plots were all dominated by BS prior to the fires. We quantified post-fire seedling density (seedlings/ha) of BS and co-occurring tree species to quantify the proportion of plots that were affected by BS regeneration failure, i.e., that shifted toward JP or deciduous species, such as paper birch (*Betula papyrifera* Marsh.) and trembling aspen (*Populus tremuloides* Michx.). The latter were abbreviated DS, or plots were categorized as non-forest vegetation types, which were abbreviated NF. We analyzed the effects of seed bank conditions, fire regime characteristics (fire severity and seasonality), and seedbed conditions on post-fire BS regeneration. We discuss our results in the context of current and future climate-induced changes in the boreal forest biome.

## 2. Material and methods

### 2.1 Study area

The study area is located in the boreal forest of eastern North America between latitudes 49°70’N-51°90’N and longitudes 73°58’W–67°68’W (Figure 1). The 21 fires under study had burned between 1995 and 2016 and are located in the black spruce-feather moss bioclimatic domain of Quebec, eastern Canada (Saucier et al. 2009). This bioclimatic domain is characterized by BS dominance with JP, balsam fir (*Abies balsamea* [L.] Mill.), paper birch and trembling aspen as companion species, depending upon site conditions and successional stage. The study area is mostly covered by glacial deposits, particularly undifferentiated till. Its climate is characterized by average annual temperatures varying between 0.05 and -2.19°C, with average annual precipitation varying between 946 and 1121 mm. The growing season is short, generally extending between May and September (Environment Canada 2024). The mean fire return interval varies along a longitudinal gradient, ranging from about 220 years near Lake Mistassini in the west to about 640 years near the Manicouagan Reservoir in the east (Bouchard et al. 2008; Bélisle et al. 2011; Couillard et al. 2022).

**Figure 1:**
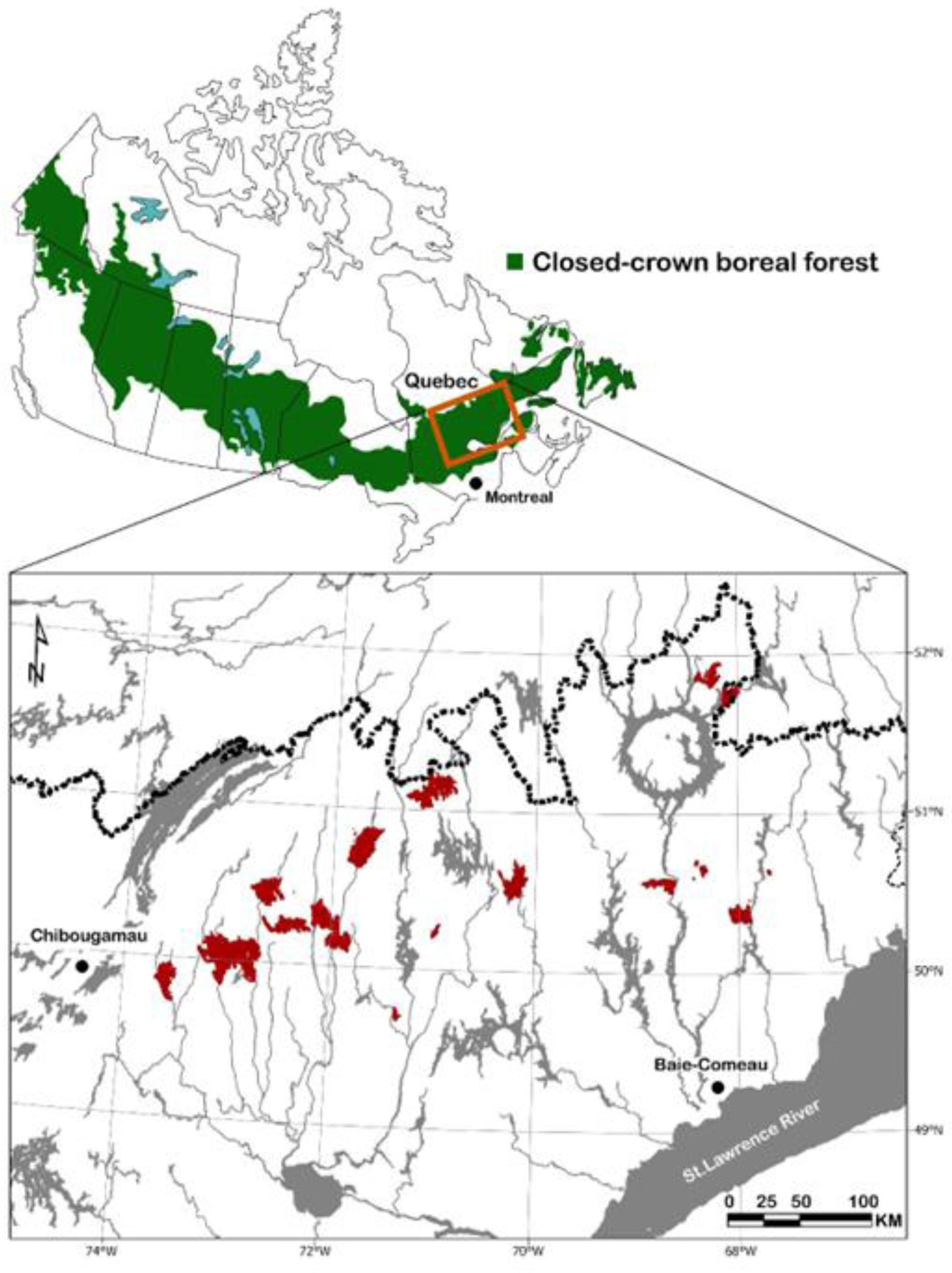
Location of the 21 studied fires (1995-2016; in red) in the closed-crown boreal forest of Quebec, eastern Canada. The hatched line corresponds to the northern boundary of the managed forest area. The closed boreal forest area is drawn according to Rowe (1972).

### 2.2 Site selection

The studied fires were selected because: (i) they were composed of mature (> 60 years) BS-dominated stands (> 50% of the basal area occupied by BS); (ii) they had burned a minimum of three years before field surveys to allow time for post-fire seedling establishment (St-Pierre et al. 1992); (iii) they had burned after 1984, allowing the acquisition of differenced Normalized Burn Ratio data (dNBR, see Danneyrolles et al. 2024); (iv) they had not been salvage logged; and (v) they were accessible by road. Across all selected fires, 536 circular plots of 400 m^2^ (r = 11.28 m) were sampled during the summers of 2017, 2018, 2019, and 2020 following a systematic sampling design. Finally, each plot was located at least 50 m away from any human disturbances (cutovers, roads), unburned stands, shrub-dominated vegetation or water bodies to avoid edge effects (Harper et al. 2005; Boucher et al. 2011) and were separated from one another by at least 100 m to minimize spatial autocorrelation.

### 2.3 Aerial seed bank conditions

To estimate aerial seed bank conditions, we evaluated the pre-fire stand basal area and age, which are good proxies of viable seed availability (Viglas et al. 2013; Splawinski et al. 2016;). In each plot, all pre-fire trees with a diameter at breast height (DBH) ≥ 1 cm were counted by species and their DBH was measured to calculate stand basal area (m^2^/ha). To determine the mean pre-fire stand age, transversal disks of five representative burned trees were collected at 0.30 m from the ground surface. Samples were dried and finely sanded in the lab to count the number of annual growth rings, and tree age was adjusted to consider sampling height (Delwaide and Filion 2010).

### 2.4 Seedbed conditions and seedling counts

The percentage cover of each understory plant species was determined in each plot using the point-intercept method (Buckland et al. 2001), along two perpendicular transect lines of 22.56 m that crossed the plot along two perpendicular axes were set in place for a total length of 45.12 m. At every 0.5 m increment along these lines, 1.5-metre-long metal rods were placed perpendicularly. The number of rods that were touched by each species across a plot was counted and divided by the total number of rods, yielding a percent cover value (Lutes et al. 2006). The percentage ground cover of understory plants was obtained, including living *Sphagnum* spp. moss and ericaceous shrubs. All living post-fire tree seedlings (height > 0.01 m) of each of the main tree species, which included BS, JP, paper birch or trembling aspen (forming 99.9% of all tree stems), were counted in 10 evenly distributed 4 m^2^ subplots. The thickness (cm) of the post-fire residual organic layer was measured at the centre of each of the 10 subplots.

### 2.5 Fire regime characteristics

Fire seasonality, which was defined as the date of fire occurrence, was assigned using the MODIS hotspots product (Giglio et al. 2018), which provides the date to which a given pixel or group of pixels burned (identified as a hotspot). As MODIS provides data only since 2001, the Canadian National Fire Database (CNFDB; CFS 2023) was used to assign a date for previous fires. With these combined data, a precise fire date could be assigned to each sample plot (S.I.1). To take into consideration the timing of snowmelt, the date of fire from each plot was adjusted to the start of the fire season, which was extracted from the software BioSim 11 (Régnière et al. 2017). The fire season was considered to start when no more snow was present around the weather stations that are nearest to a given plot for three consecutive days or when the temperature is at least 12°C at midday (CFS 1984). The beginning of the fire season was designated as Julian Day 1, and the Julian date for each fire was determined accordingly.

Fire severity (*sensu* Keeley 2009) in each plot was determined using the differenced normalized burn ratio (dNBR) that was calculated as per Key and Benson (2006). Google Earth Engine was used to extract cloud-free pre- and post-fire images that were taken from the Landsat collection 2 imagery. The resulting dNBR values were stored in 900 m^2^ pixels. A weighted average dNBR value was calculated when a 400 m^2^ plot was covered by more than one pixel. The Composite Burn Index (CBI), ranging between 0 (unburned) and 3 (highest severity) (Key and Benson 2006), was then calculated from the equations of Danneyrolles et al. (2024), which were specifically developed for the study region.

### 2.6 Statistical analysis

#### 2.6.1 Evaluation of post-fire regeneration and changes in forest composition

Post-fire regeneration level was assessed in two different ways. First, based on post-fire recovery studies (Boiffin and Munson 2013; Perrault-Hébert et al. 2017) and forestry regeneration standards (FEC 2023), we quantified the absolute post-fire regeneration (PFR^A^) of each plot by classifying global density of BS and JP seedlings (seedlings/ha) into three broad categories: regeneration failure (< 1750 seedlings/ha); low regeneration (between 1750 and 8000 seedlings/ha); or adequate regeneration (> 8000 seedlings/ha). JP was included in the analysis, because it is an important fire-adapted coniferous companion species to BS in the eastern Canadian boreal forest (Lavoie and Sirois 1998; see S.I.2). Second, to better account for the effect of pre-fire stem density, the relative post-fire regeneration index (PFR_R_) was calculated for each plot as the ratio of post-fire BS and JP seedling density to their pre-fire stem density. This index reflects the maintenance of tree species density within a plot, where a PFR_R_ ≥ 1.00 indicates post-fire regeneration density equal to or exceeding pre-fire stem density levels, independent of the regeneration failure threshold that was mentioned above.

To determine whether there was a change in successional trajectory after fire (i.e., pre-fire mature BS-dominated stands were replaced by JP-, DS- or NF-dominated stands), the proportion of each plot that was occupied by seedlings of each main tree species or group of species (BS, JP or DS) after fire was calculated. A given species or group of species was considered dominant if it represented 50% or more of the total number of post-fire seedlings.

#### 2.6.2 Environmental variables influencing post-fire BS regeneration

Based on the reproductive ecology of BS, as documented in the scientific literature, we selected measurable and interpretable environmental variables that could influence BS post-fire regeneration (Perrault-Hébert et al. 2017; Boucher et al. 2020). These variables provide insight into (i) seed bank condition, (ii) fire regime characteristics, and (iii) seedbed conditions (Table 1). To avoid collinearity, we used the variance inflation factor (VIF) function of the *car* package, and only the explanatory variables with a VIF < 2 were retained (Dormann et al. 2013). A total of six readily measurable and interpretable variables were thus retained from which explanatory models of post-fire BS regeneration were constructed (Table 1; S.I.3).

**Table 1:**
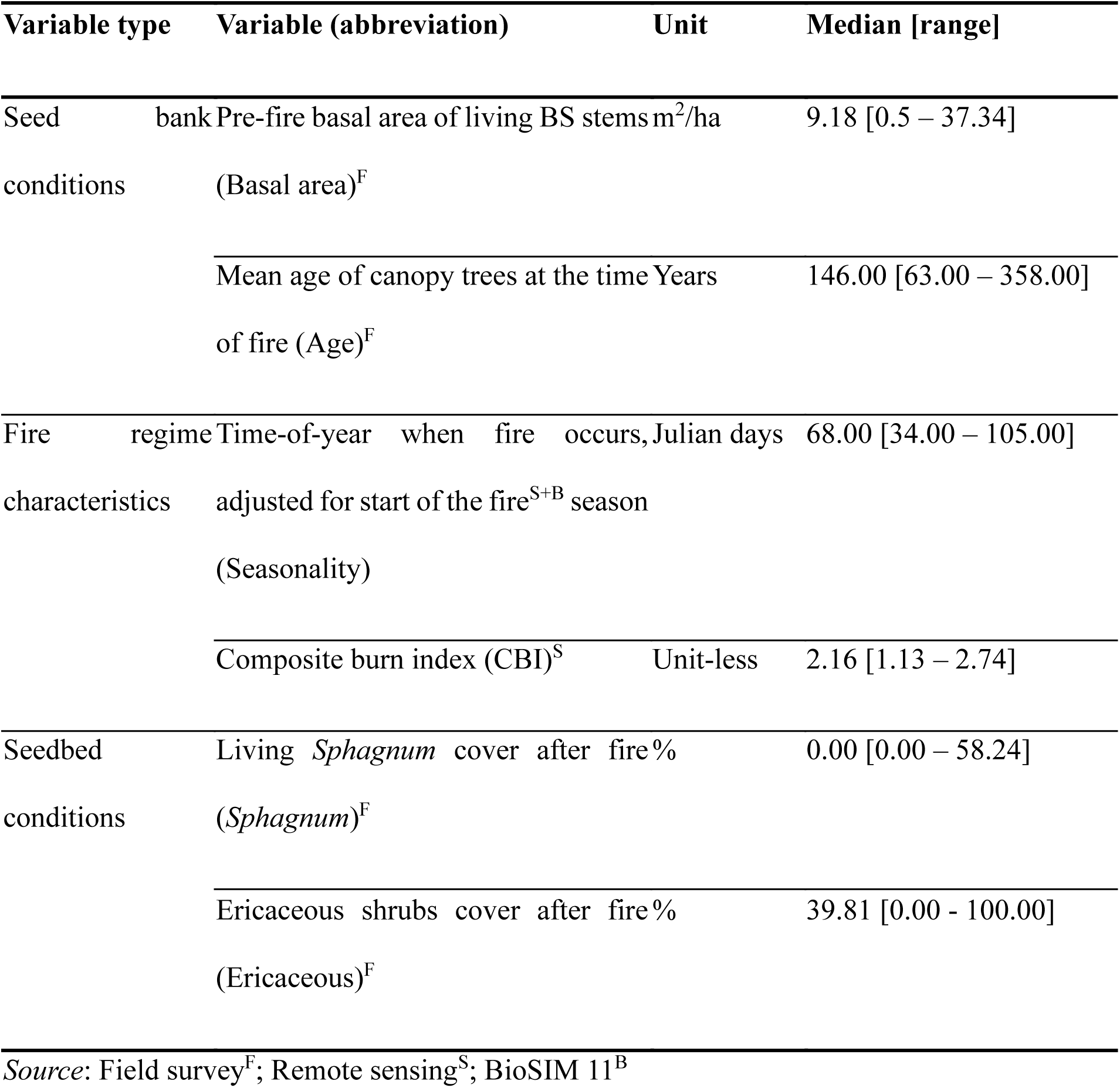
Description of the six explanatory environmental variables selected to model post-fire BS seedling density.

Generalized linear mixed-effect models (GLMM) with a negative binomial distribution (*glmmTMB* package in R; Brook et al. 2017) were used to model BS seedling density as a function of the different combinations of the explanatory variables. A total of 51 candidate models were constructed by combining the six explanatory variables. Normality prerequisites of residuals were met (*DHARMa* package in R; Hartig 2022). Due to the large study area, a random effect corresponding to the identifier of each fire was included to account for unmeasured differences between fires. The Moran I statistic for spatial autocorrelation was used to assess whether residuals of the candidate models still exhibited spatial autocorrelation (Table 2) (*spdep* package in R; Bivand et al. 2013).

**Table 2:** The four top-ranking models that were retained for model averaging among the 51 candidate models predicting BS seedling density, as assessed with the corrected Aikaike’s information criterion (AICc). The estimated parameters are provided, including the intercept (K), the AICc values, the difference in AIC (ΔAIC), the AIC weight (AICwt) and the cumulative AIC weight (CUMwt). The Moran I statistic for spatial autocorrelation (Moran I) and its associated *p*-value are also provided.

| Model rank | Variables | K | AICc | $\Delta AIC$ | AICwt | CUMwt | Moran's I<br>($p$-value) |
| --- | --- | --- | --- | --- | --- | --- | --- |
| 1 | Basal area + CBI + Seasonality<br>+ *Sphagnum* | 7 | 4167.17 | 0.00 | 0.45 | 0.45 | 0.014<br>(0.16) |
| 2 | Basal area + CBI + *Sphagnum* | 6 | 4168.13 | 0.96 | 0.28 | 0.73 | 0.014<br>(0.16) |
| 3 | Basal area + CBI + Seasonality<br>+ *Sphagnum* + Age | 8 | 4168.99 | 1.82 | 0.18 | 0.91 | 0.013<br>(0.18) |
| 4 | Basal area + CBI + Seasonality<br>+ *Sphagnum* + Age +<br>Ericaceous | 9 | 4170.44 | 3.27 | 0.09 | 1.00 | 0.015<br>(0.15) |

The candidate models were ranked according to their AIC values (Burnham and Anderson 2002) using the *AICmodavg* package (Mazerolle 2023). Since the top-ranking models had AIC weights (AICwt) < 0.95, model averaging was performed. Average parameter estimates and associated unconditional standard errors and unconditional 90% confidence intervals were calculated for the top four ranking models, for which the sum of AICwt values exceeded 0.95.

Predictive models of BS seedling density were made at the population level according to the values of the explanatory variables retained during model averaging. For each explanatory variable, their values were classified according to the three PFR_A_ classes and the mean value of each group was compared using ANOVAs and Tukey’s tests (*TukeyHSD* function of the *stats* package in R). Following the same principle, the values of each explanatory variable were separated according to post-fire species dominance. Given that the data were not normally distributed, a Kruskal-Wallis test (non-parametric one-way ANOVA) was used to determine whether the group means differed (*Kruskal.test* function of the *stats* package in R). All statistics were calculated using R software (version 4.3.2; R Development Core Team 2023).

## 3. Results

### 3.1 Evaluation of post-fire regeneration and changes in forest composition

Overall, according to our classification, 33.4% of the plots experienced regeneration failure (< 1750 seedlings/ha), 41.0% exhibited low regeneration (1750 to 8000 seedlings/ha), and 25.6% had adequate regeneration (> 8000 seedlings/ha). Among all sample plots, 72.6% had a pre-fire stem density above the regeneration failure seedling density (≥ 1750 stems/ha) and 27.4% had a pre-fire stem density below the regeneration failure seedling density (< 1750 stems/ha; Figure 2a). Most regeneration failures were observed in plots with pre-fire stem density above the regeneration failure threshold (i.e., >1750 stems/ha; Figure 2b). Among these plots which were above the regeneration failure threshold (< 1750 stems/ha) before fire, 41.1% showed a reduction in density after fire (PFR_R_ < 1.0); the mode of the PFR_R_ was 0.45, and the median PFR_R_ was 1.03 (Figure 2c). Among the plots with a pre-fire stem density below the regeneration failure threshold (< 1750 stems/ha), 21.1% showed a reduction in density after fire (PFR_R_ < 1.0); the mode and the median of the PFR_R_ were 1.47, and 3.20, respectively (Figure 2c).

**Figure 2:**
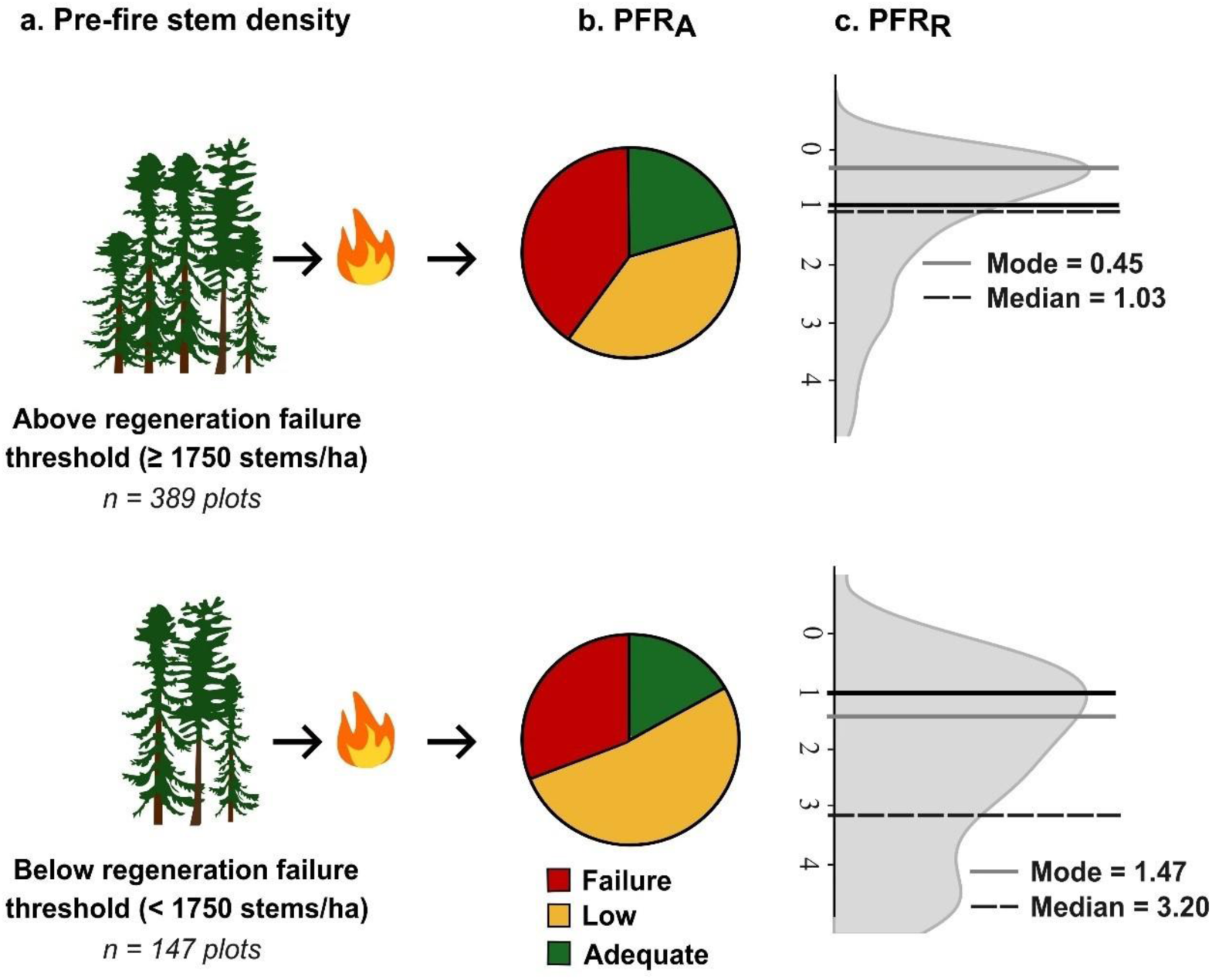
Post-fire regeneration in plots whose pre-fire stem density (a) was either above regeneration failure threshold (≥ 1750 stems/ha; top row) or below regeneration failure threshold (< 1750 stems/ha; bottom row). (b) The absolute post-fire regeneration classes (PFR_A_) distinguish between plots having an adequate seedling density (> 8000 seedlings/ha), plots with low seedling density (1750 to 8000 seedlings/ha), and plots that experienced post-fire regeneration failure due to an insufficient density of seedlings (< 1750 seedlings/ha). (c) Frequency distribution of the relative post-fire regeneration index (PFR_R_), calculated as the ratio of each plot’s post-fire seedling density (BS + JP) to its pre-fire stem density. For each distribution, the continuous black line indicates where PFR_R_ = 1, the continuous grey line indicates the mode of the PFR_R_, and the hatched black line indicates the median of the PFR_R_. For clarity, PFR_R_ values > 5 are not shown.

Considering the entire data set (n = 536 plots; Figure 3a), 81.0% of the plots remained dominated by BS after fire, 12.3% became dominated by JP, 2.6% by DS, and 4.1% transitioned to NF vegetation types (Figure 3b). Among the plots that remained dominated by BS, 30.9% showed regeneration failure, 44.7% had low regeneration, and 24.4% had adequate regeneration. Post-fire BS-dominated plots also experienced a reduction in density, with 41.4% showing a PFR_R_ < 1.00 (Figure 3c, d). Among the plots in which dominance shifted to JP, more than half (54.6%) experienced regeneration failure. Yet, when compared to pre-fire density, only a slight loss of density was observed (Figure 3c, d). Of the few (2.6%) plots that had recorded a change in dominance to DS stands, half experienced regeneration failure, while the remainder exhibited low regeneration. Accordingly, most plots showed a much lower density after fire (Figure 3c, d). Finally, 4.1% of all plots exhibited complete regeneration failure and had transitioned to NF types (Figure 3c, d).

**Figure 3:**
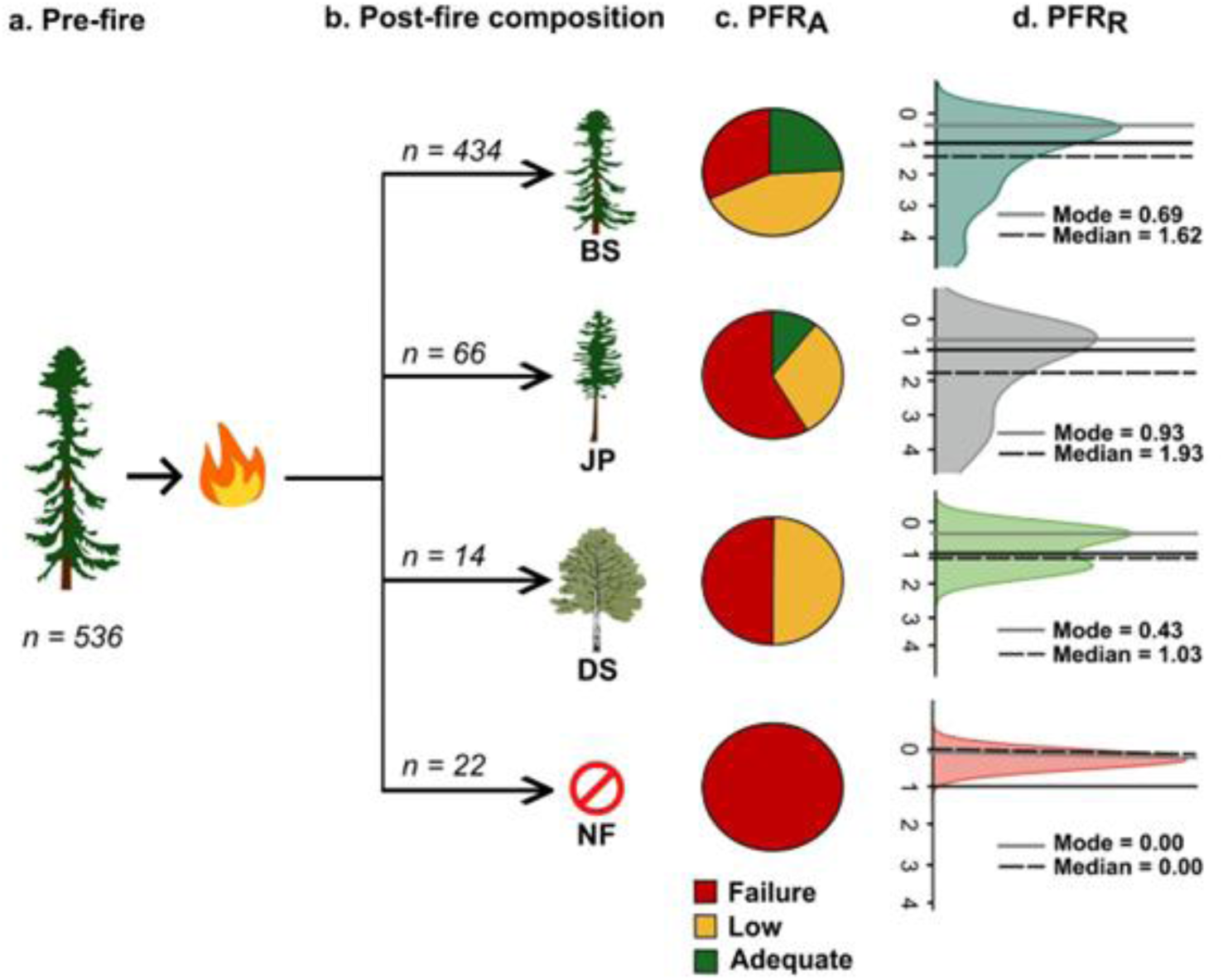
Post-fire regeneration according to post-fire stand composition. (a) Number of plots dominated by BS before fire. (b) Number of plots whose post-fire composition either stayed dominated by black spruce (BS; top row), shifted to a dominance by jack pine (JP; 2nd row), shifted to a dominance by deciduous species (DS; 3rd row), or converted to non-forest (NF; bottom row) due to regeneration failure. (c) The absolute post-fire regeneration classes (PFR_A_) distinguish plots having an adequate BS and JP seedling density (> 8000 seedlings/ha), plots with low seedling density (1750 to 8000 seedlings/ha), and plots that experienced post-fire regeneration failure due to an insufficient density of seedlings (< 1750 seedlings/ha). (d) Frequency distribution of the relative post-fire regeneration index (PFR_R_), calculated as the ratio of each plot’s post-fire seedling density to its pre-fire stem density. For each distribution, the continuous black line indicates where PFR_R_ = 1, the continuous grey line indicates the mode of the PFR_R_, and the hatched black line indicates the median of the PFR_R_. For clarity, PFR_R_ values > 5 are not shown.

### 3.2 Environmental variables influencing post-fire BS regeneration

Among the four top-ranking models (CumWT of 1.00; Table 2) predicting post-fire density of BS seedlings, all of the candidates included *Sphagnum*, Basal area and CBI, while three also included seasonality. These four environmental variables were significant (within 90% confidence intervals (CI)) but the other two (Age and Ericaceous) were not (S.I.4). The top-ranking model (AICWT = 0.45) explained 60% of the variation in BS seedling density (conditional *R^2^* = 0.60 and marginal *R^2^* = 0.43).

BS seedling density increased with *Sphagnum* cover (average coefficient = 0.092, *p*-value < 0.001; S.I.4) and, thus, plots showing adequate regeneration had a higher percentage of *Sphagnum* cover than did plots that failed to regenerate, which were mostly devoid of moss (Figure 4a). A similar positive relationship was observed for Basal area (average coefficient = 0.048, *p*-value < 0.001; S.I.4), with adequate regeneration more frequently occurring in plots with higher pre-fire basal areas (Figure 4b). Seasonality was also positively correlated (average coefficient = 0.013, *p*-value < 0.1; S.I.4) with BS seedling density, although the correlation was weaker than with the other variables. Regeneration failures were more frequent in fires that occurred earlier in the season, whereas low regeneration and adequate regeneration levels were more frequent later in the season (Figure 4c). The Composite Burn Index (CBI) was, however, negatively correlated (average coefficient = -0.61, *p*-value < 0.001; S.I.4) with post-fire seedling density. Therefore, the plots that had regenerated adequately had a lower CBI than those that had failed. Higher CBI values tended to favour a transition to DS stands (Figure 4d).

**Figure 4:**
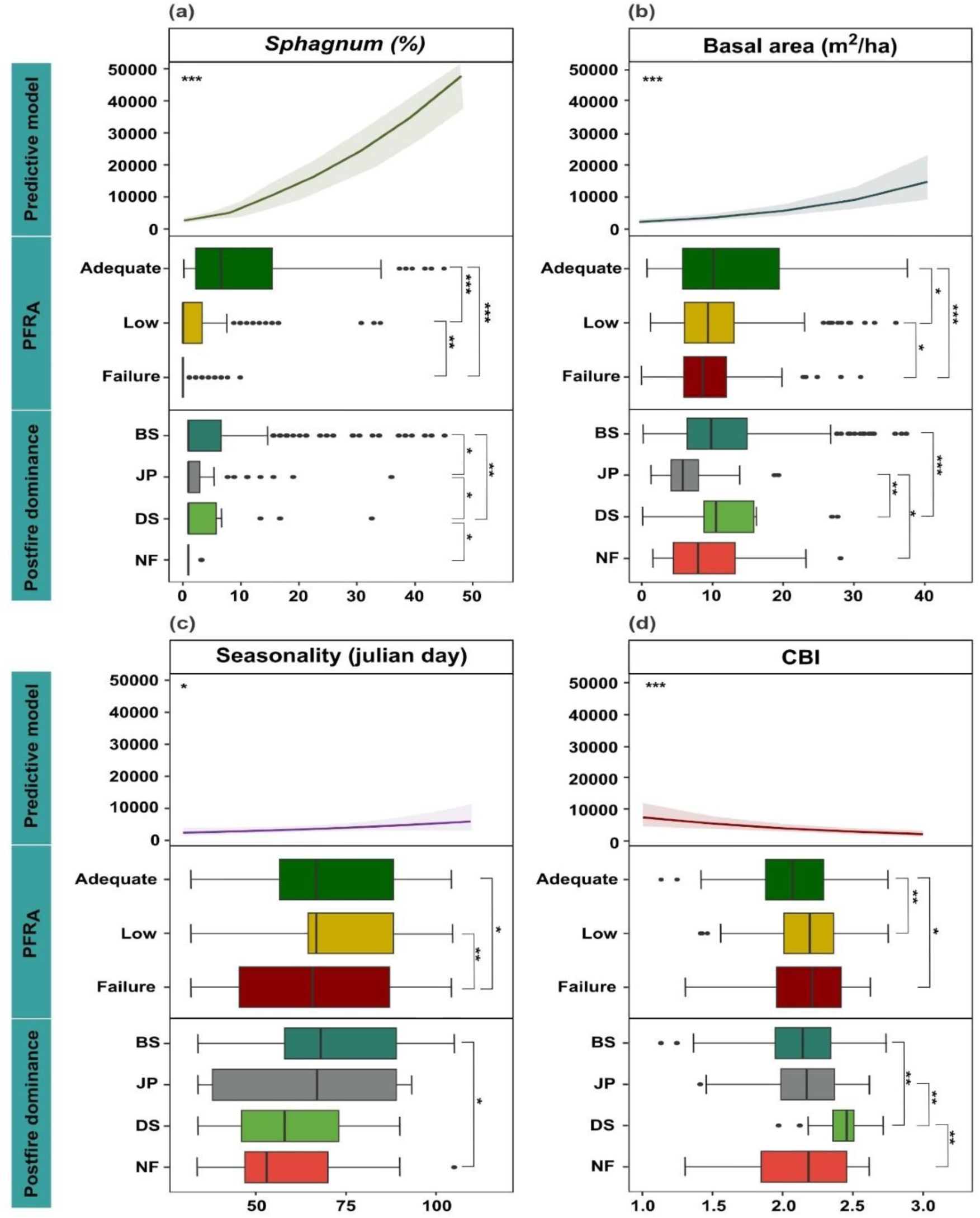
Predictive models of post-fire black spruce (BS) regeneration (seedlings/ha) and frequency distribution of predictive variable values by PFR_A_ classes and post-fire vegetation dominance for the following predictive variables: (a) post-fire *Sphagnum* cover, (b) Pre-fire basal area, (c) Fire seasonality, and (d) Composite Burn Index (CBI). The 90% confidence interval is represented by the shaded area on the predictive model graphs. The asterisks represent the level of marginal significance (*** = 0.001, ** = 0.01, * = 0.1). BS, black spruce; JP, jack pine; DS, deciduous species; NF, non-forest.

## 4. Discussion

The increase in fire activity induced by climate warming is raising serious concerns regarding the resilience of boreal forests and their capacity to provide ecosystem services (Gauthier et al. 2015; Hartmann et al. 2022). To our knowledge, this is the first study to demonstrate, at the subcontinental scale, the negative influence of high-severity canopy fires on post-fire recovery of mature BS-dominated boreal forests. Our study identified several keys spatially explicit predictors of BS regeneration, which are crucial for monitoring post-fire resilience and for informing restoration efforts should broad-scale regeneration failure be observed (D’Amato et al. 2023; Thiffault et al. 2023).

A large fraction of the stands that we studied experienced post-fire regeneration failure. While immature BS-dominated stands are at high risk of post-fire regeneration failure (Girard et al. 2009; Perrault-Hébert et al. 2017; Hart et al. 2019), it is generally assumed that mature BS-dominated stands are resilient to fire (Viereck and Johnston 1990; Splawinski et al. 2019a). Immature trees have a small viable seed bank that limits their sexual regeneration capacity (Brown and Johnstone 2012; Viglas et al. 2013; Whitman et al. 2019). Our results showed that stands with a higher pre-fire basal area of BS had a higher density of seedlings after fire. This is likely because BS basal area is positively related to the size of the aerial seed bank (Greene and Johnson 1998; Splawinski et al. 2019b). Although most of the stands under study were dominated by trees that were older than 100 years (mean age ± SD: 159 ± 61 years; S.I.5), an age at which BS seed banks normally are fully developed, one-third of the studied stands failed to regenerate. Outbreaks of eastern spruce budworm (*Choristoneura fumiferana*), which primarily affect such old forests, may have occurred prior to fire, thereby reducing the BS seed bank by feeding on spruce flowers, cones, and seeds (Schooley 1980; Simard and Payette 2005). An extensive analysis of aerial surveys of insect defoliation during the 10 years preceding each studied fire, however, revealed that less than 1% of the plots had been affected by this insect defoliator (Bouchard and Auger 2014). Rather, our results would suggest that high-severity crown fires, which can deplete the BS aerial seed bank through intense combustion (Zasada et al. 1979; Splawinski et al. 2019b; Day et al. 2022; Reid et al. 2023), represent the main factor limiting post-fire seedling establishment and subsequent regeneration failure that was observed in our study region. Our results are also consistent with those of Perrault-Hébert et al. (2017), whose study was the first to document the negative effect of high severity fires on BS regeneration, based upon dNBR assessments.

Unburned patches of living *Sphagnum* moss can act as refugia that help enhance the post-fire resilience of BS. Our analysis clearly indicates that post-fire regeneration success is strongly associated with living *Sphagnum* spp. cover, given that 34.9% of all seedlings (n = 17 154 seedlings) were located on this substrate. After fire, residual living patches of *Sphagnum* have high moisture contents and, as such, provide a good seedbed, considering that blackened organic matter tends to be dry following fire due to its lower albedo (Greene et al. 2004; Veilleux-Nolin and Payette 2012; Boiffin and Munson 2013; Perrault-Hébert et al. 2017). Yet, it is generally assumed that exposed mineral soil is the most frequent and favourable seedbed for BS seedling establishment in the boreal forest of North America, particularly in the west (Thomas and Wien 1985; Charron and Greene 2002; Greene et al. 2004; Wang and Kemball 2005). This substrate was quite rare in our 21 burns, given the substantial thickness of the residual organic matter that was observed in our study sites, i.e., 17.9 ± 9.7 cm (S.I.6). Consequently, it is unlikely that fire could produce enough exposed mineral soil to support adequate post-fire regeneration of BS (Boiffin and Munson 2013). Indeed, eastern North American boreal forests tend to accumulate more organic matter than do western North American boreal forests, due in part to the wetter climate and the generally lower fire frequency that is observed in the east (Bergeron et al. 2004; Carcaillet et al. 2006). Although living *Sphagnum* is a very good microsite for BS post-fire establishment, it can hamper BS growth because thicker organic matter accumulations cause the water table to rise, which can lower nutrient availability in the organic soil layer (Munson and Timmer 1989; Fenton et al. 2006; Lavoie et al. 2007; Pacé et al. 2018). Although high densities of BS seedlings were observed in small patches of unburned *Sphagnum*, this response is likely to lead to self-thinning mortality over the next few decades, exacerbating the problem of poor regeneration. While BS may persist on burned sites with *Sphagnum* moss, which will allow the return of regeneration and therefore ensure the resilience of the forest, the suboptimal conditions for growth and space could slow down stand productivity in the long-term.

The occurrence of fire early in the fire season was another important characteristic of the fire regime affecting post-fire recovery. Our results corroborate previous studies demonstrating that BS regeneration is generally lower following spring fires compared to those occurring later in the fire season (Girard et al. 2009; Le Page et al. 2010; Veilleux-Nolin and Payette 2012). A widely discussed hypothesis suggests that spring fires may lead to poor BS seedling establishment by reducing the density of exposed bare mineral soil, given that waterlogged or frozen soil during snowmelt causes the fire to burn the organic horizon only superficially (Heinselman 1981; Miyanishi and Johnson 2002; Veilleux-Nolin and Payette 2012). In addition, spring fires occur when the previous season’s leaves and herbs are dead and physiological processes are re-initiating, coinciding with a short period of high conifer leaf flammability. This flammability is due to a low water-to-carbon ratio in the needles, a phenomenon known as “spring dip” (Jolly et al. 2014). However, the leafing out of deciduous vegetation, or “greenup,” including the leaf flush of canopy-forming trees, has an even more influential effect on landscape flammability. This period, when forests produce very high-density fuels, is known as the “spring window” by fire managers, given that it makes forests very sensitive to fire ignition and conducive to fire spread (Parisien et al. 2023). Moreover, climatic conditions are particularly conducive to fire during hot and dry springs (Heinselman 1981; Bonan and Shuggart 1989; Johnson 1992; Wang 2024). Globally, these conditions have the potential to induce severe fires, which are expected to damage the BS aerial seed bank. Drawing on insights from the eastern Canadian boreal forests, we argue that BS regeneration is more limited by high burn severity at the canopy level, which reduces seed survival, than by restricted access to mineral soil seedbeds (Charron and Greene 2002; Veilleux-Nolin and Payette 2012).

Unlike the frequently observed decline in BS density following a fire, post-fire compositional changes were relatively minor. BS-dominated stands remained dominated by BS after fire, whereas JP was the only tree species that substantially benefited from fires to increase its dominance. This expansion of JP over BS could be explained by several ecological traits favouring the species in a context of increasing fire activity. First, JP reaches maximum seed production earlier (5-10 years; Viereck 1983) and its cones have a higher degree of serotiny than do those of BS, which confers upon JP an advantage when fire is either severe or occurs in young stands. Second, the seeds of JP are considerably larger than those of BS (Burns and Honkala 1990). Consequently, JP seedlings have a faster and longer radicle elongation, providing critical access to water resources that are located deeper in the underlying mineral soil (Thomas and Wein 1985; Johnstone and Chapin 2006). The post-fire increase in JP dominance in our study is consistent with other studies that were conducted in the same region (see Boiffin and Munson 2013; Baltzer et al. 2021). Transitions to DS stands were rare (< 3%), probably because the fertility, drainage, and climate of our study area are suboptimal for the growth of paper birch and trembling aspen (Boucher et al. 2014). Finally, a significant part of the studied stands (4.1%) experienced a transition to NF vegetation types. The occurrence of this fire-induced alternative stable state (forest to non-forest) is of concern due to the predicted increase in fire frequency and severity in the coming decades (Coogan et al. 2019; Wang 2024).

## 5. Conclusion

Our work highlights the key drivers of post-fire BS regeneration in the boreal forest of eastern North America. High-severity fires have been identified as strongly reducing BS regeneration potential, presumably by damaging the aerial seed bank in mature stands. The presence of living *Sphagnum* moss after fire is another important factor that provides refugia for seedling establishment, enabling the regeneration of BS and maintaining its resilience, especially in areas with limited exposed mineral soil. Our results further indicate that post-fire seedling density was significantly reduced compared to pre-fire stem density. In contrast, these fires did not lead to major changes in successional trajectories and forest composition. JP has increased its dominance and may be considered an option for reinforcing resilience if the restoration of sites experiencing regeneration failure is being considered by forest managers. In North America, an increase in spring wildfires has already been observed (Jain et al. 2017; Hanes et al. 2019; Burton 2023; Wang 2024), and climate warming will likely exacerbate this phenomenon (Flannigan et al. 2013). Moreover, increased fire severity due to a higher number of days with extreme fire weather is also expected in the coming decades (Whitman et al. 2018, Augustin et al. 2022, Boulanger et al. 2025). Management interventions such as fuel reduction treatments or prescribed burning have been proposed to reduce fire severity risks (see Davis et al. 2023), but these are impractical given the vast areas that would need to be treated. Alternatively, forest managers should set up broad-scale monitoring programs to track post-fire regeneration. This would allow for the planning of reactive restoration activities such as planting or seeding BS, JP, or other species potentially able to cope with the new fire regime characteristics (Boulanger et al. 2025). In light of our results, the predicted trends in boreal fire regimes should raise serious concerns about the resilience of BS-dominated boreal forests and their capacity to provide ecosystem services over the long-term.

## Acknowledgements

This project was funded by la Direction de la recherche forestière du ministère des Ressources Naturelles et des Forêts du Gouvernement du Québec (project #142332132 to Y. Boucher), a NSERC RDC (O. Valeria, R. Fournier and Y. Boucher), a NSERC Alliance (Y. Bergeron and Y. Boucher), and the Observatoire régional de recherche sur la forêt boréale de l’Université du Québec à Chicoutimi (Y. Boucher). We thank representatives of Domtar, Produits Forestiers Arbec, and Société Sylvicole Mistassini for providing lodging for this study’s four years of sampling. We would particularly thank H. Tremblay, M. Perrault-Hébert, S. Marcouiller, B. Bour and several interns for contributing to summer field sampling. We also thank I. Auger, F. Rousseu, D. Bonfils and H. Dorion for their assistance with statistical analysis. W.F.J. Parsons edited the English text.

## Authors contributions

**Stelsa Fortin:** conceptualization, data curation, formal analysis, investigation, methodology, visualization, writing – original draft writing, review and editing. **Yan Boucher:** funding acquisitions, supervision, conceptualization, data curation, formal analysis, investigation, methodology, visualization, writing – original draft writing – review and editing. **Yves Bergeron:** supervision, writing – review and editing. **Martin Simard:** formal analysis, investigation, methodology, visualization, writing– review and editing. **Dominique Arsenault:** writing– review and editing. **Hugo Asselin:** writing– review and editing. **Martin Barrette:** writing – review and editing. **Victor Danneyrolles:** visualization, writing – original draft, review and editing. **Sylvie Gauthier:** writing– review and editing. **Francois Girard:** writing– review and editing. **Martin Girardin:** writing– review and editing. **Marc-André Parisien:** writing– review and editing. **Nelson Thiffault:** writing– review and editing. **Osvaldo Valeria:** funding acquisition, writing– review and editing.

