## Supplemental files for "High-severity fires undermine resilience of black spruce-dominated boreal forests in eastern North America"

**Supplementary files**

**S.I.1:** Date of burning of the sampling plots according to fire identifier

| Fire identifier | Year of fire | Date of fire |
|---|---|---|
| **1995_1** | 1995 | 11 August |
| **2002_1** | 2002 | 10 June |
| **2005_1** | 2005 | 5-6 June |
| **2005_2** | | 2-3 and 5 June |
| **2005_3** | | 3 June |
| **2005_4** | | 4-5 June |
| **2005_5** | | 4 June |
| **2005_6** | | 2 June |
| **2007_1** | 2007 | 22 June |
| **2007_2** | | 18-19 June |
| **2007_3** | | 16-18 June |
| **2007_4** | | 13-14, 16 and 19 June |
| **2007_5** | | 16 June |
| **2009_1** | 2009 | 24-25 June |

| | | |
|---|---|---|
| **2010_1** | 2010 | 22-25 June |
| **2010_2** | | 19 May |
| **2011_1** | 2011 | 15 June |
| **2013_1** | 2013 | 8-9 July |
| **2013_2** | | 9-10 July |
| **2014_1** | 2014 | 7 June |
| **2016_1** | 2016 | 12 June |

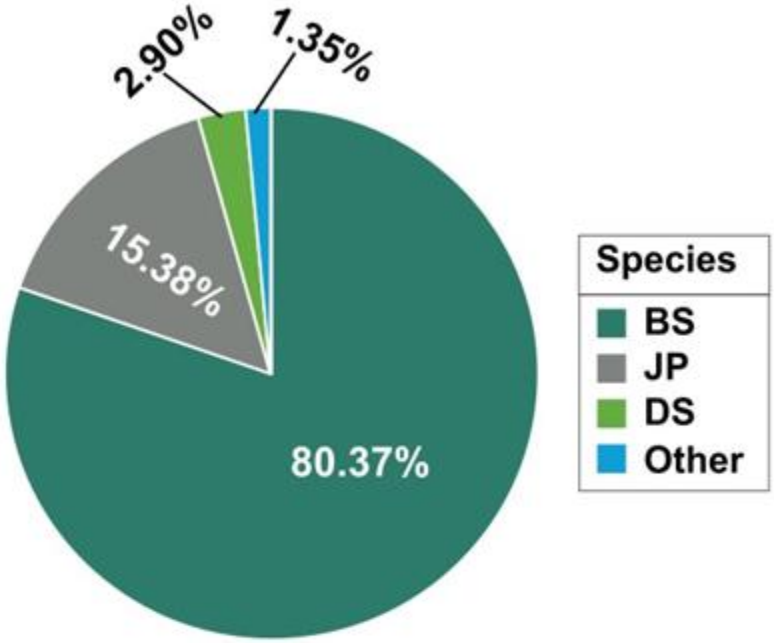


**S.I.2:** Percentage of seedlings of each species in the 536 plots (n = 17 154 seedlings). BS corresponds to black spruce (*Picea mariana*), JP to jack pine (*Pinus banksiana*), DS to deciduous species, which include paper birch (*Betula papyrifera*) and trembling aspen (*Populus tremuloides*) and Other includes balsam fir (*Abies balsamea* [L.] Mill.) and eastern larch (*Larix laricina* [Du Roi] K. Koch).

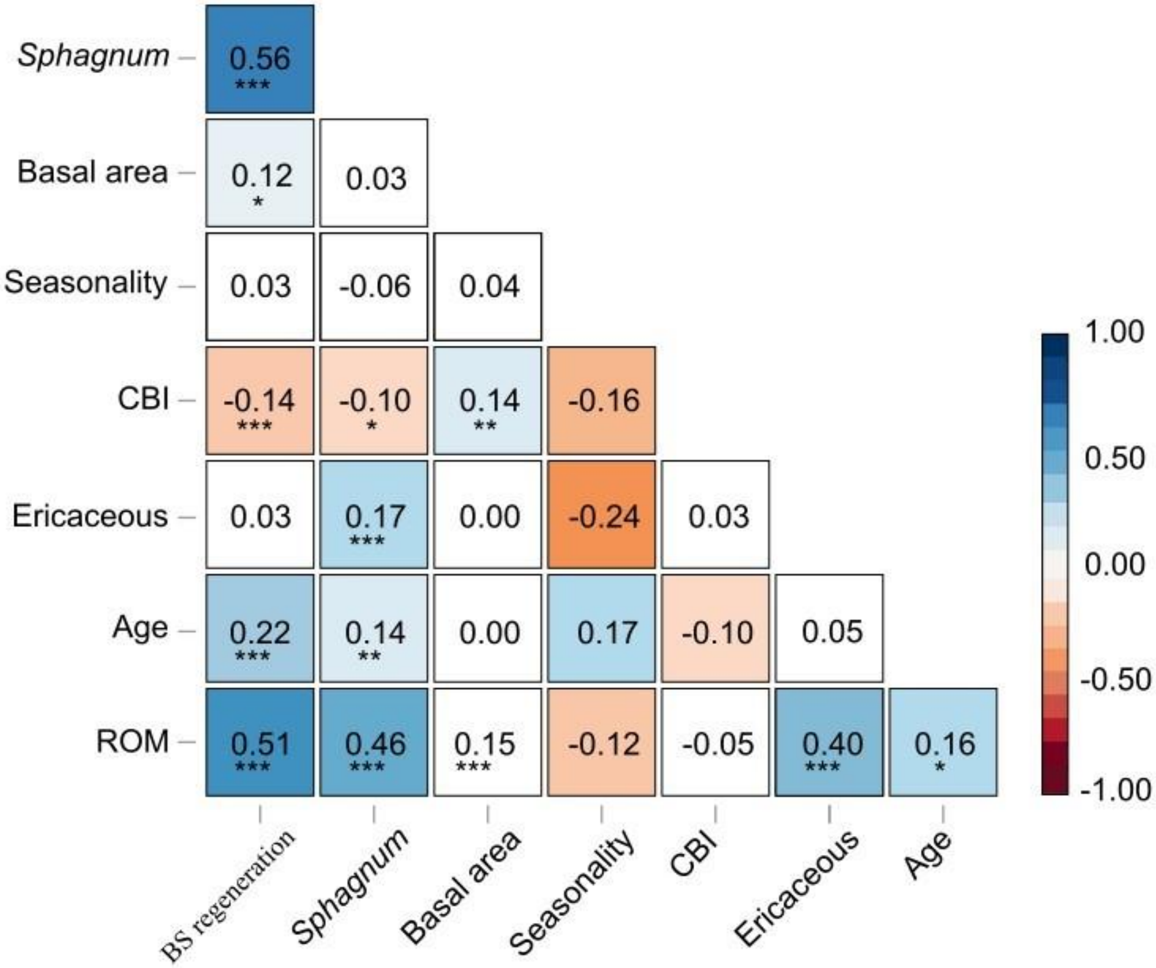


**S.I.3:** Correlation matrix between the response variable (BS seedling density) and the explanatory environmental variables using the corrected Pearson's correlation for spatial autocorrelation (modified *t*-test of the *SpatialPack* package in R; Vallejos et al. 2020). Asterisks represent the level of marginal significance given by Tukey's tests (*** = 0.001, ** = 0.01, * = 0.05).

**S.I.4:** Average coefficients and 90% confidence intervals (CI) for each variable of the four top-ranking models (AICWT > 0.90). The variables in bold are significant, given that their CI values exclude 0.

| Variables | Average coefficient | Confidence interval [lower, upper limit] |
|---|---|---|
| **Basal area** | **0.0478** | **[0.0374, 0.0582]** |
| ***Sphagnum*** | **0.0921** | **[0.0797, 0.1045]** |
| **CBI** | **-0.6053** | **[-0.8891, -0.3215]** |
| **Seasonality** | **0.0125** | **[0.0004, 0.0245]** |
| Age | -0.0002 | [-0.0018, 0.0012] |
| Ericaceous | 0.0017 | [-0.0024, 0.0059] |

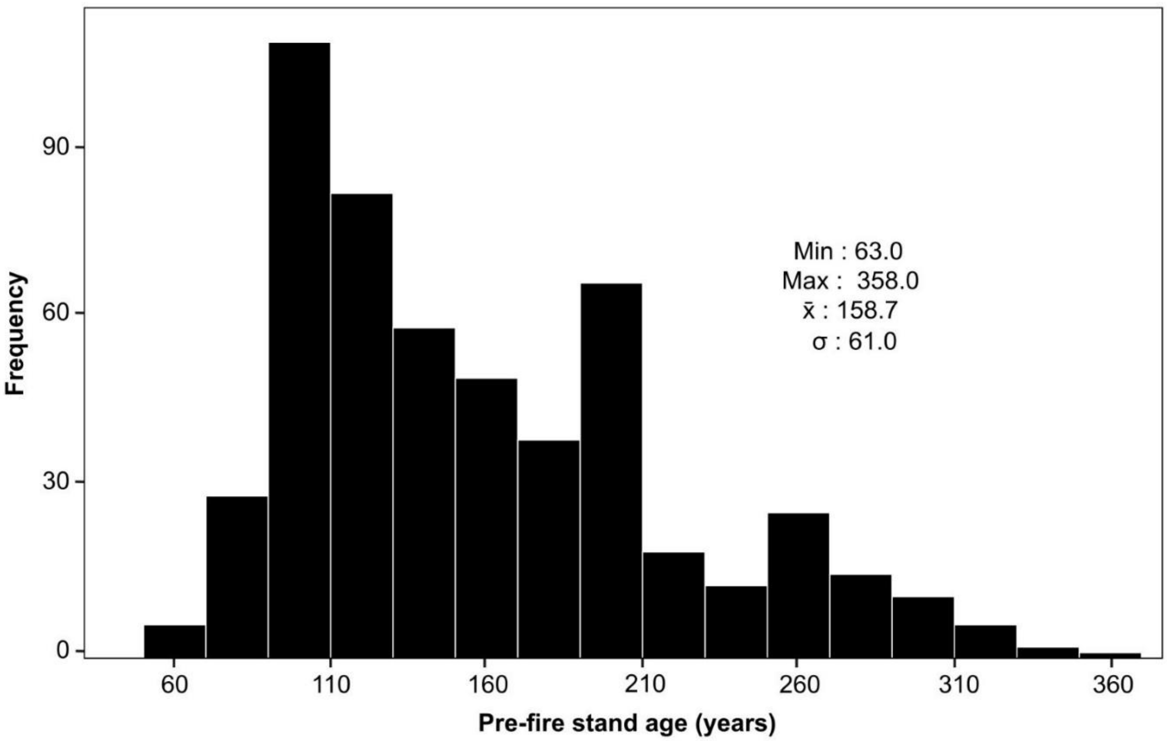


**S.I.5:** Frequency distribution and descriptive statistics of pre-fire stand age. The minimum, maximum, mean ($\bar{x}$) and standard deviation ($\sigma$) values are indicated.

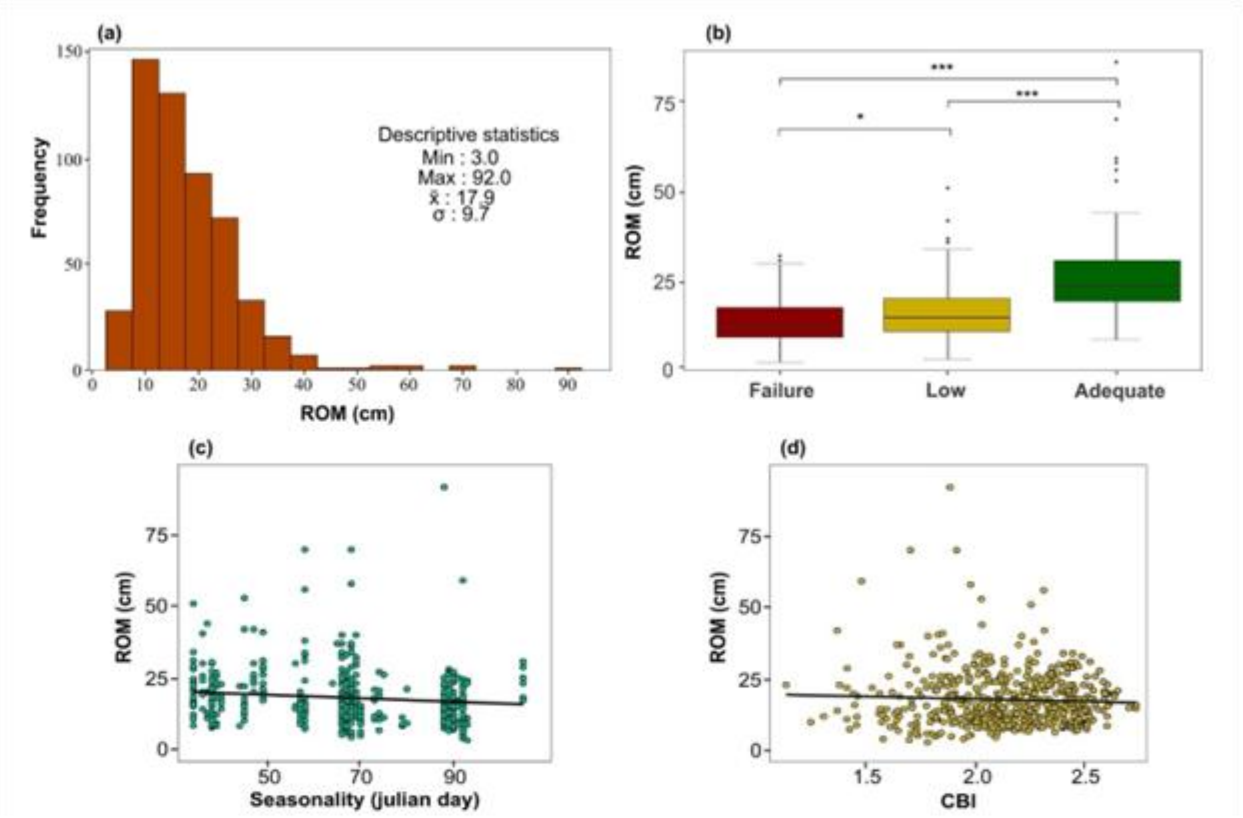


**S.I.6:** Frequency distribution and descriptive statistics of the residual organic matter (ROM) thickness. The minimum, maximum, mean ($\bar{x}$) and standard deviation ($\sigma$) values are indicated. (a). Mean ROM thickness for each of the three absolute post-fire regeneration classes ($PFR_A$) (b). Asterisks represent the level of marginal significance given by Tukey's tests (*** = 0.001, ** = 0.01, * = 0.05). ROM thickness as a function of fire seasonality (c) and the composite burn index (CBI) (d).
